# RAD50 is a potential biomarker for breast cancer diagnosis and prognosis

**DOI:** 10.1101/2024.09.07.611821

**Authors:** Kunwer S. Chhatwal, Hengrui Liu

## Abstract

**BACKGROUND:** RAD50 is one of the most critical genes in DNA double-strand break processing, which can lead to a single 3’ strand of DNA overhang and is potentially involved in forcing incomplete DNA repair. This research study aims to investigate the role of RAD50 in breast cancer diagnosis and prognosis.

**METHODS:** Breast cancer mRNA expression data was collected from TCGA and the difference between cancer and non-cancer in gene expression of RAD50 was analyzed. The survival association of RAD50 was also analyzed.

**RESULTS:** RAD50 expression is significantly lower in cancer than in normal tissue. High expression of RAD50 is associated with worse survival.

**Conclusion:** RAD50 is a potential biomarker for breast cancer diagnosis and prognosis

## 1. Introduction

Cancer is a multifaceted disease with a significant global impact, affecting both humans and other organisms. While the incidence of cancer is higher in individuals over 50 years of age, the disease’s complexity and unpredictability allow it to manifest across all age groups. According to the American Cancer Society, approximately 13% of women are expected to develop breast cancer during their lifetime, a statistic that underscores the urgent need for improved prognostic and diagnostic tools, particularly given the high fatality rate associated with the disease[1]. Breast cancer has been extensively studied in previous research, with numerous studies focusing on its molecular mechanisms, risk factors, and potential treatment strategies[2-7]. Biomarkers have become a key focus in cancer research, as they provide critical insights for early detection and disease progression. However, the current repertoire of biomarkers remains inadequate, necessitating further exploration to enhance cancer diagnosis and prognosis.

This study aims to contribute to this body of research by identifying novel biomarkers for cancer, with a specific focus on breast cancer. The gene set utilized in this research is derived from the Gene Ontology Biological Process (GOBP) database and is associated with DNA double-strand break (DSB) processing. Improper repair of DNA double-strand breaks can result in significant genomic instability, a known contributor to carcinogenesis, particularly in breast cancer[8]. By investigating the role of genes involved in this pathway, this study seeks to identify a potential biomarker that may aid in the early detection and prognosis of breast cancer, particularly in relation to the BRCA gene.

Although this research focuses on breast cancer, the implications of identifying a biomarker associated with DNA damage repair mechanisms could extend to other cancer types, given the universal importance of maintaining genomic integrity across various forms of the disease. Bioinformatics has been widely utilized in the study of human diseases, with numerous databases playing a critical role in advancing research[9-11]. These databases allow researchers to analyze large-scale genomic, proteomic, and transcriptomic data, facilitating the identification of disease-associated genes, pathways, and biomarkers. By integrating various data sources, bioinformatics enables a more comprehensive understanding of disease mechanisms and contributes to the development of personalized medicine approaches. The application of bioinformatics tools and databases has proven especially valuable in fields like cancer research, where complex genetic and molecular alterations require sophisticated analytical techniques to uncover meaningful patterns and insights.

In this study, we applied bioinformatics approaches to explore potential genetic biomarkers for breast cancer. By leveraging publicly available datasets and conducting a comprehensive analysis of gene expression profiles, we aimed to identify novel biomarkers that could aid in the early diagnosis and prognosis of breast cancer. The findings from this research contribute to a broader understanding of cancer biology by highlighting key genetic factors involved in tumor development and progression. The identification of such biomarkers holds promise for improving diagnostic accuracy and prognostic strategies, not only in breast cancer but potentially across the broader cancer spectrum. These results may pave the way for more personalized treatment options, ultimately enhancing patient outcomes.

## 2. Methods

### 2.1. Data Acquisition

The data utilized for this study were obtained from The Cancer Genome Atlas (TCGA) database, focusing on breast cancer (BRCA) patients. The RAD50 gene expression data were collected for both tumor and normal breast tissue samples. The dataset consisted of RNA sequencing (RNA-seq) results presented as log2-transformed RSEM (RNA-Seq by Expectation-Maximization) normalized counts. Clinical survival data, including overall survival (OS) times, were also extracted from TCGA to correlate RAD50 expression with patient outcomes.

### 2.2. Gene Expression Analysis

RAD50 expression was analyzed by comparing tumor and normal breast tissues using boxplots. The expression levels between the two groups were statistically evaluated, and the false discovery rate (FDR) was calculated to assess the significance of the expression difference. An FDR threshold of 0.05 was used to determine statistical significance.

### 2.3. Survival Analysis

To assess the prognostic significance of RAD50 expression in breast cancer, a Kaplan-Meier survival analysis was conducted. Patients were stratified into two groups based on median RAD50 expression levels: those with higher expression and those with lower expression. The overall survival (OS) probabilities were calculated for both groups, and the survival curves were compared using the log-rank test. A P-value less than 0.05 was considered statistically significant. Additionally, the hazard ratio (HR) was computed to determine the relative risk associated with RAD50 expression.

## 3. Results

### 3.1. Differential Expression of RAD50 in Breast Tumor vs. Normal Tissues

RAD50 expression was significantly elevated in breast tumor tissues compared to normal tissues (Figure 1). The boxplot demonstrates that tumor samples exhibit a higher median RAD50 expression level (log2 RSEM) than normal samples, with an FDR of 1.2e-02, indicating statistical significance. This finding suggests that RAD50 may play a role in breast cancer development and could be a potential marker for tumor presence.

**Figure 1.**
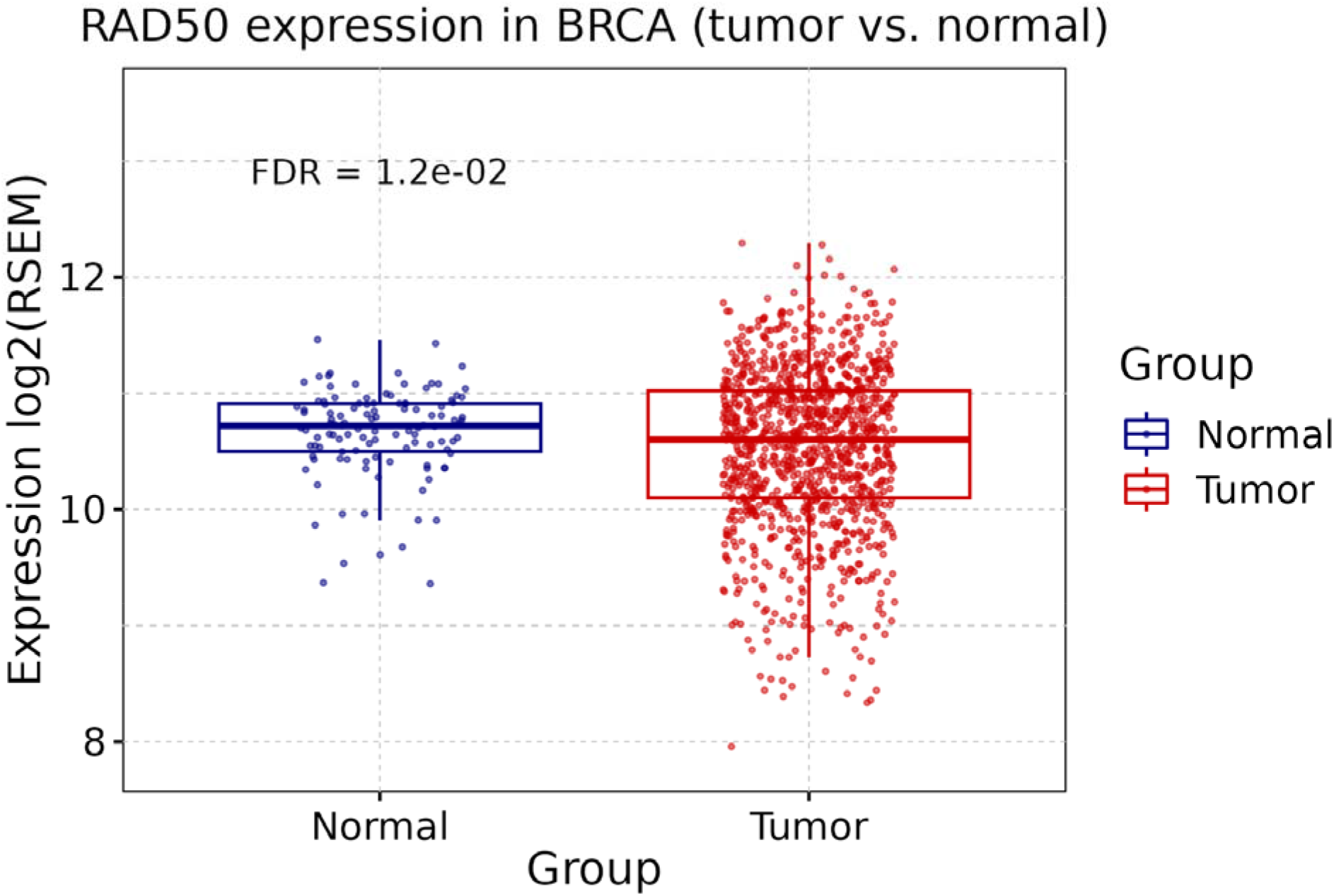
Expression of RAD50 in Breast tumor compare to in normal tissues. FDR: false discovery rate.

### 3.4. Survival Analysis of RAD50 Expression in Breast Cancer

The Kaplan-Meier survival analysis revealed a significant association between higher RAD50 expression and poorer overall survival in breast cancer patients (Figure 2). Patients with elevated RAD50 expression (n=547) had a notably lower OS probability compared to those with lower RAD50 expression (n=547). The log-rank P-value was 0.046, confirming the statistical significance of the survival difference. The hazard ratio for patients with high RAD50 expression was 1.39, indicating that increased RAD50 expression is associated with a 39% higher risk of mortality. These results suggest that RAD50 may be a significant prognostic biomarker in breast cancer, with its overexpression correlating with worse patient outcomes.

**Figure 2.**
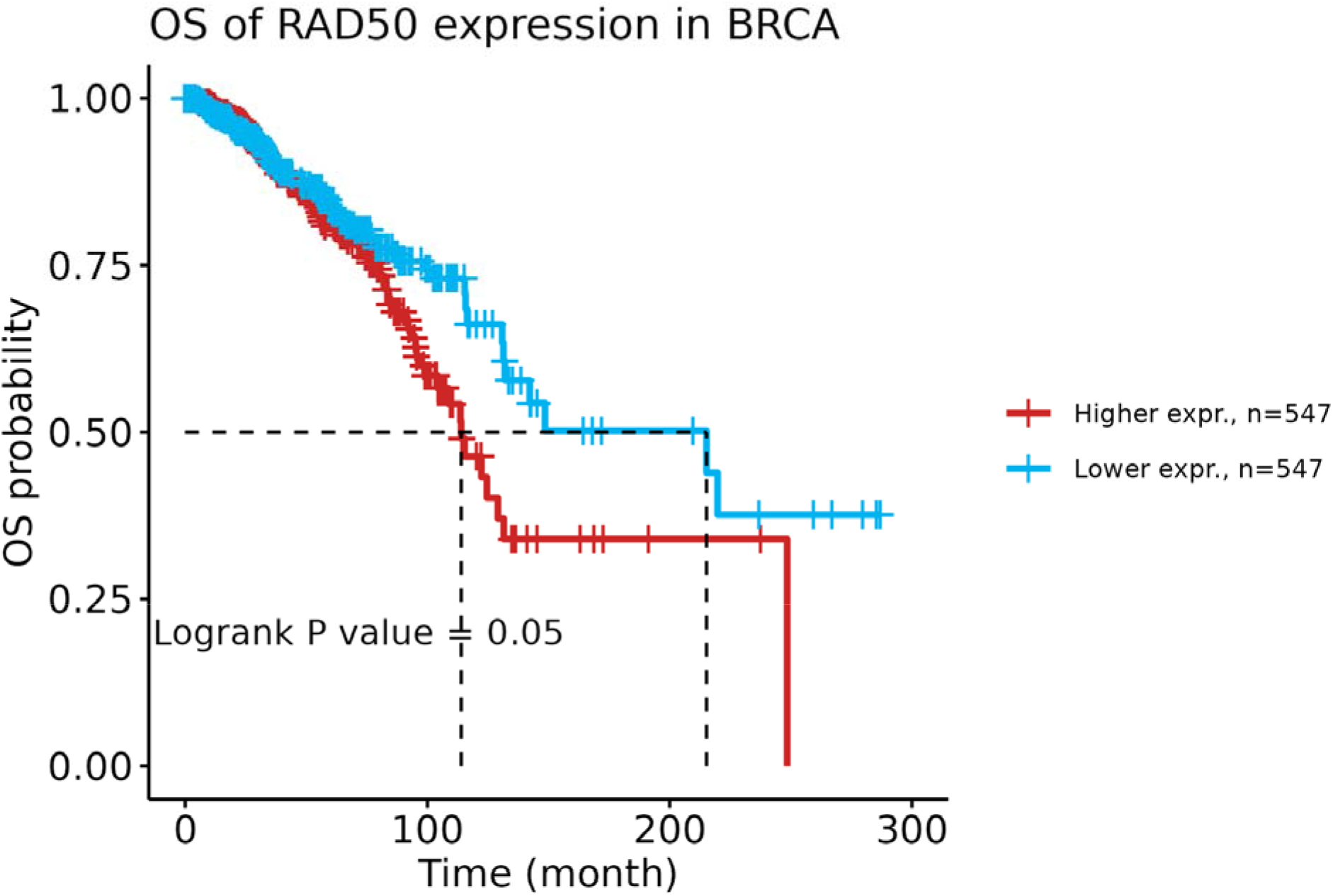
Kaplan-Meier survival curve illustrating the overall survival (OS) probability of breast cancer (BRCA) patients stratified by RAD50 expression levels. The cohort is divided into two groups: patients with higher RAD50 expression (red line, n=547) and those with lower RAD50 expression (blue line, n=547).

## 4. Discussion

This study applied TCGA data to study cancer genes and biomarkers as in many previously studies[12-28]. Based on the results of this study, it is clear that RAD50 expression is significantly associated with the prognosis of breast cancer (BRCA), particularly in the context of DNA double-strand break (DSB) repair mechanisms. A previous study reported RAD51, another gene from the same family, as a cancer biomarker[21]. The overexpression of RAD50 in tumor tissues compared to normal tissues, combined with its correlation with poorer overall survival, suggests that RAD50 may be a critical player in the DNA repair processes that contribute to breast cancer progression.

While investigating the role of DNA double-strand break (DSB) repair mechanisms in breast cancer, this study reviewed the literature to identify relevant studies that highlight the importance of these mechanisms. A particularly notable study reported that C1QBP promotes homologous recombination by stabilizing MRE11 and regulating the assembly and activation of the MRE11/RAD50/NBS1 complex[29]. This complex is crucial in the detection and repair of DSBs, which, when improperly regulated, can contribute to genomic instability and the development of cancer. The current study, focusing specifically on human breast cancer (BRCA), provides insights that have direct clinical relevance. In particular, the identification of RAD50 as a potential biomarker for breast cancer prognosis represents a novel and significant contribution to the field. Although the role of DSB repair mechanisms has been extensively explored in various contexts, the direct association between RAD50 expression and breast cancer patient survival has not been sufficiently studied.

Kaplan-Meier survival analysis conducted in this study revealed that patients with higher RAD50 expression exhibit significantly worse overall survival outcomes compared to those with lower RAD50 expression. This finding suggests that RAD50 overexpression may aggravate the consequences of faulty DNA repair, leading to more aggressive cancer phenotypes and worse clinical outcomes. Therefore, RAD50’s role in the MRE11/RAD50/NBS1 complex and its association with homologous recombination repair mechanisms underscore its potential as a prognostic biomarker for breast cancer. Further exploration of RAD50’s functional role in DNA repair could provide additional avenues for therapeutic intervention and patient stratification in breast cancer treatment. Furthermore, while Ashworth’s study provided a foundation for understanding DSB repair mechanisms, it did not specifically focus on RAD50’s prognostic value in breast cancer. This research adds to the existing body of knowledge by demonstrating the potential clinical relevance of RAD50 expression in human breast cancer. As such, RAD50 could serve as a biomarker to stratify breast cancer patients based on their risk levels, potentially guiding treatment decisions and improving patient outcomes.

## 5. Conclusion

While previous research on DSB repair mechanisms has advanced our understanding of cancer biology, this study expands on this by providing human-specific insights, particularly regarding the role of RAD50 in breast cancer prognosis. These findings may contribute to improved diagnostic and prognostic tools, enhancing personalized approaches to breast cancer treatment.

## 6. Declearation

### 6.1. Ethical Approval and Consent to participate

Not applicable.

### 6.2. Availability of supporting data

The source of the raw data was provided in the paper and the raw analysis data of this study are provided by the corresponding author with a reasonable request.

### 6.3. Competing interests

There is no conflict of interest.

### 6.4. Authors’ contributions

All the Analyses were done by Kunwer S. Chhatwal. Kunwer S. Chhatwal drafted the manuscript and Hengrui Liu significantly modified it. Hengrui Liu supervised the project and direct this study.

### 6.5. Funding

This study received no funding.

## 6.6. Acknowledgments

The author thanks the support of Weifen Chen, Zongxiong Liu, Bryan Liu, and Yaqi Yang.

## 6.7. Other information

This study is conducted during the Lumiere academic Project where Kunwer S. Chhatwal was a student and Hengrui Liu was the academic research project mentor.

## Notes

### Competing Interest Statement

The authors have declared no competing interest.

